# Integration of Odor-Induced Activity of Kenyon Cells in an Electrotonically Compact *Drosophila* Mushroom Body Output Neuron (MBON)

**DOI:** 10.1101/847244

**Authors:** Omar A. Hafez, Benjamin Escribano, Jan Pielage, Ernst Niebur

## Abstract

The formation of an ecologically useful lasting memory requires that the brain has an accurate internal representation of the surrounding environment. In addition, it must have the ability to integrate a variety of different sensory stimuli and associate them with rewarding and aversive behavioral outcomes. Over the previous years, a number of studies have dissected the anatomy and elucidated some of the working principles of the *Drosophila* mushroom body (MB), the fly’s center for learning and memory. As a consequence, we now have a functional understanding of where and how in the MB sensory stimuli converge and are associated. However, the molecular and cellular dynamics at the critical synaptic intersection for this process, the Kenyon cell-mushroom body output neuron (KC-MBON) synapse, are largely unknown. Here, we introduce a first approach to understand this integration process and the physiological changes occurring at the KC-MBON synapse during Kenyon cell (KC) activation. We use the published connectome of the *Drosophila* MB to construct a functional computational model of the MBON-α3 dendritic structure. We simulate synaptic input by individual KC-MBON synapses by current injections into precisely (*μm*) identified local dendritic sections, and the input from a model population of KCs representing an odor by a spatially distributed cluster of current injections. By recording the effect of the simulated current injections on the membrane potential of the neuron, we show that the MBON-α3 is electrotonically compact. This suggests that odor-induced MBON activity is likely governed by input strength while the positions of KC input synapses are largely irrelevant.

## 1 Introduction

For living organisms, interactions with the environment strongly depend on their capability to form an internal representation of the surrounding world. Storing information about potential rewards and dangers requires a constant update of such representations and it is crucial for survival. This allows the organism to adapt its future behavior with the goal of increasing rewarding interactions with the environment. In the brain, information of different stimuli can be associated through learning and then memorised over long periods of time. Recall of such associations will then lead to an adapted behavioral output.

Here, we use the *Drosophila melanogaster* mushroom body (MB) as a model to study the cellular basis of learning and memory. The MB is required for acquisition and recall of associative olfactory memories[12, 24, 31, 32, 37, 51]. Among the advantages of this structure is that the number of neurons in the MB is relatively small and that, in Drosophila, many of these neurons are genetically accessible through a large variety of tools that allow visualisation and neuronal manipulation of neuronal subpopulations [13, 16, 23, 25, 27, 30, 36, 39, 44, 48]. The MB consists of about 2000 Kenyon cells (KCs) per hemisphere that receive stochastic innervation by projection neurons (PNs) from the antennal lobes [29, 46]. The antennal lobes, in turn, receive direct input from the olfactory receptor neurons (ORNs) of the antennae [52]. Stochastic innervation of KCs therefore generates a random and sparse representation of odors in the MB [7, 21, 50]. The somata of all KCs are clustered in the posterior brain and project their axons into the MB calyx, the horizontal and vertical lobes of the MB [11, 26, 28, 47]. KCs then innervate 22 mushroom body output neurons (MBONs) per hemisphere in a highly compartmentalised manner [3]. Current data supports a balance model, in which the net output activity of the MBONs defines the behavior of the fly when it encounters a previously conditioned odor [2, 3, 9].

Information regarding reward or punishment of an odor is provided by dopaminergic neurons (DANs) [6] that innervate the KC-MBON synapse that also respect the compartmentalised nature of the MB [10]. Coincident activation of a KC and local release of dopamine is thought to selectively depress KC-MBON synapses resulting in modified MBON activity [4, 35, 40, 42] and, as a consequence, altered behavior of the fly [19]. Therefore, the output behavior to the conditioned odor will be changed [49]. Despite the accumulated knowledge of the structure of the MB and the functional model of its influence on fly behavior, the molecular and cellular processes at the KC-MBON synapse inducing and maintaining decreased activation of the MBON remain largely unknown.

Importantly, the recent complete electron-microscopic (EM) reconstruction of the *Drosophila* MB [45] and the entire fly brain [53] provided precise information of the anatomical structure of KC, DAN and MBONs and their synaptic interactions. In addition, we recently developed a novel tool to selectively mark, and manipulate the activity of, individual KCs that store associative aversive olfactory long-term memory [43]. This tool enables us to directly access KCs and to evaluate synaptic connectivity changes to the MBONs that are relevant to modulate behavioral changes.

In the present study, we combine our knowledge of KC-based storage of long-term memories [43] with the precise anatomical data provided by EM-studies [14, 45] to determine the nature of KC-MBON interaction. As a starting point we use the MBON-α3 neuron which receives 12,770 input synapses from KCs. We perform computational simulations to determine the relevance of individual synaptic input along the dendritic tree to provide an estimate regarding the number of synapses and KCs necessary to evoke MBON activation. Our study provides first evidence that MBON-α3 is electrotonically compact and demonstrates that few KCs may be sufficient to evoke reliable MBON activation.

## 2 Results

In this study we make use of the known dendritic structure of the MBON-α3 to generate a computational model of the effect of input KC synapses on the membrane potential in this neuron. Based on electrophysiological data, Cassenaer and Laurent [8] have suggested that Drosophila MBON neurons are electrotonically compact. One purpose of our computational study is to determine to what extent this is the case for one specific neuron, MBON-α3. Given the observed unipolar morphology of an MBON, we assumed passive (linear) cable-theory to model the neuron’s functionality. In general, it is not yet understood to what extent nonlinear currents occur in the dendritic tree of MBONs. Therefore, we will test to what extent the neuron is functional in their complete absence. We consider this, on the one hand, as a necessary baseline condition that can be extended if either experimental evidence becomes available that requires revision of this hypothesis, or if the assumption of passive dendrites is demonstrably not compatible with the functionality of this neuron. On the other hand, it is the simplest assumption that can be made for this structure, and if the computational results show that the neuron can fulfill its function under this assumption, an argument can be made that no more complex mechanisms are required. This simple model is then preferable over more complex ones by the principle of “Occam’s Razor.”

If the neuron is electrotonically compact, small current injections made at different locations along its dendritic tree will have identical effects on the membrane potential at the neuron’s proximal neurite. We, furthermore, hypothesize that a small number of injected currents with the characteristics of typical excitatory synaptic inputs, representing the typical input pattern received from KCs, would be sufficient to excite the MBON by depolarizing its proximal neurite past the threshold for firing action potentials. This hypothesis is motivated by the fact that, to compensate for the small number of active synapses for any one odor, the synapses between KCs and the MBONs are particularly strong, with an average excitatory post-synaptic potential (EPSP) of 1.58mV [8].

Using the NEURON environment [20], we simulated such small current injections at different locations along the MBON-α3 dendritic tree. To anticipate our results, our data provide first evidence that MBON-α3 is electrotonically compact. Furthermore it provides insight into the integration of learned olfactory information at MBON dendritic trees.

We first (Section 2.1) show examples of excitations at randomly selected synapses. Subsequently, we systematically excite selected segments (Section 2.3) and patterns of segments (Section 2.2). Note that stimulation currents in Section 2.1 have a duration of 100ms (and a correspondingly low current to generate EPSPs compatible with experimental observations). This is meant only as an illustration and to probe the neuron’s behavior with a simple stimulation protocol that allows us to compare the shape of the voltage excursion over a longer time course (Fig. 1H below). Data in Section 2.3 and 2.2 are obtained with a stimulation pattern where each dendritic segment (= model synapse) is excited by a square current pulse of *5ms* duration. These current injections are a simple approximation to fast synaptic activation currents, like those resulting from glutamatergic α-amino-3-hydroxy-5-methyl-4-isoxazolepropionic acid receptor (AMPA-R) type synapses. To guide the reader, Table 1 gives an overview of all Results figures.

**Figure 1:**
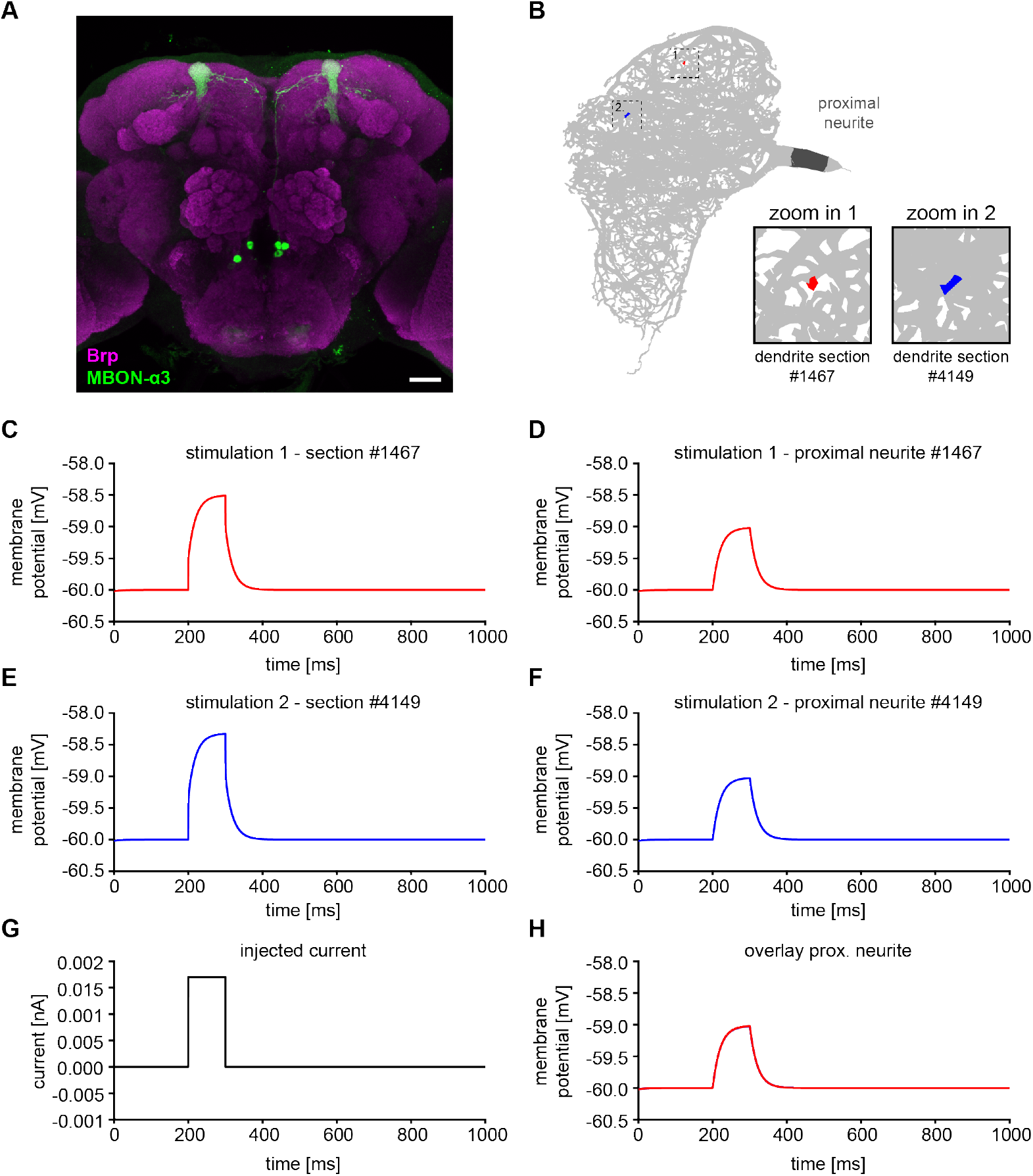
Constructing a computational model of MBON-α3. (A) Immunohistological staining of MBON-α3 (green) and Bruchpilot (Brp) as a marker for neuronal active zones (magenta). See Section 4.2 for details. (B) Structure of the Drosophila MBON-α3 dendritic tree visualised by NEURON. Morphology of the dendritic arborisations as characterised by Takemura *et al.* [45]. The morphology of the soma and axon are not known. Magnified boxes represent stimulated dendritic segment #1467 (red) and #4149 (blue). The proximal neurite at the base of the dendrite is darkened for visualization. A square-pulse current (see G) was injected into either dendritic section #1467 (C) or #4149 (E), resulting in a depolarization of the membrane potential by approximately +1.6*mV* at the location of current injection. The effect of these current pulses on the proximal neurite is shown in (D) for segment #1467 and in (F) for #4149. In both cases, it resulted in membrane potential depolarisations of about 1.0*mV*. (G) Protocol of injected current with an amplitude of 1.7*pA* for 100*ms*. (H) Overlay of membrane potentials in proximal neurite resulting from the two current injections (from D and F) shows nearly identical voltage excursions.

**Table 1:** Overview of Results figures.

| Fig. | I[nA] | Pulse length [ms] | # synapses | Recording site | max V[mV] |
| --- | --- | --- | --- | --- | --- |
| 1C,E | 0.0017 | 100 | 1 | dendrite | -58.5 (C), -58.3 (E) |
| 1D,F | 0.0017 | 100 | 1 | proximal neurite | -59 |
| 3B,C | 0.02 | 5 | 1 | dendrite | $\approx [-55, -45]$ |
| 3D,E | 0.02 | 5 | 1 | proximal neurite | $\approx [-58, -57]$ |
| 2B,C | 0.02 | 5 | 25 | proximal neurite | $\approx [3, 6]$ |

### 2.1 Time course of voltage excursions in dendrite and proximal neurite

Simulated current was injected into one randomly selected dendritic section (#1467; Figure 1B, C), and the resulting change in membrane potential at the proximal neurite was recorded. Before current injection, the simulation was run for 200*ms* simulated time without input to allow the system to equilibrate, after which current with an amplitude of 1.7*pA* was injected for^1^ 100ms (Figure 1G). This was sufficient to depolarize the dendritic membrane voltage in this segment by +1.5mV (Figure 1C), the approximate size of a typical EPSP for a KC – MBON synapse [8]. The simulation lasted for 1,000ms in total duration. This was repeated at a different random location on the dendrite tree (#4149; Figure 1B, E) to determine whether the location of current injection influenced the change in membrane potential at the proximal neurite (Figure 1D,F). Figure 1H shows that the voltage traces in the proximal neurite from the two excitations overlap nearly perfectly, suggesting that the location of synaptic input has very little influence on their effect on proximal neurite excitation. This is an indication that the neuron is electrotonically compact. In the following Section we will investigate how general this result is by systematically exciting a large variety of segments in the dendritic tree.

### 2.2 Distributions of voltage excursions in dendrite and proximal neurite for distributed activation patterns of synapses

It was previously established that about 5% of all Kenyon cells are activated upon stimulation by one odor [7]. Former studies have also shown that a subset of engram cells in ≈ 500 *αβ* surface neurons (a subclass of KCs) are required for memory recall [22, 43]. Therefore we hypothesized that excitation of 5% of all *αβ* surface neurons would represent a memory-relevant and physiologically realistic odor induced activation of an MBON. We reduced the network to one KC-MBON synapse per KC for a first simplified approach. We therefore studied the depolarization at the proximal neurite following simultaneous current pulse stimulation of 25 segments distributed over the MBON’s dendritic tree. Synaptic activation was simulated in each of these segments by injected current pulses with an amplitude of 0.02nA and duration of 5ms (same as in Section 2.3 below).

Figure 2B shows an overlay of all voltage traces for 200 trials, each with a different set of randomly selected stimulated segments. Supplementary Figure S1B,C shows the corresponding histogram of maximal voltage excursions in the proximal neurite. Peak voltages were all positive (min: 2.82mV, max: 6.18mV, average: 4.23mV). The fact that this distribution is wider than that of the individual EPSPs at the proximal neurite (Figure 3E) may indicate that nonlinear interactions between currents become important. However, all combined EPSPs are far above spiking threshold, with a substantial “safety factor,” so functionally this may be of limited relevance.

**Figure 2:**
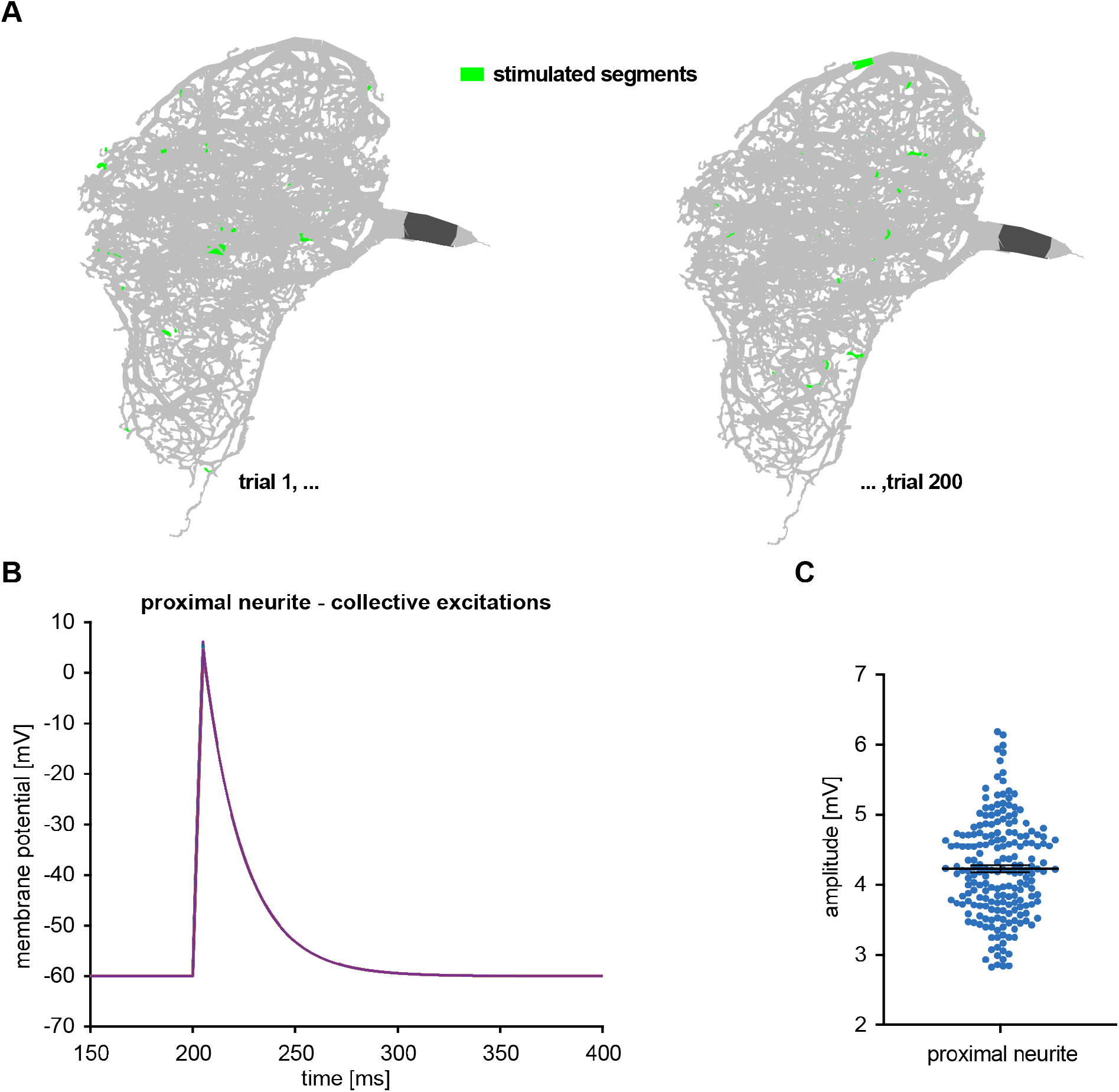
Simultaneous current injections of multiple sections in the MBON-α3 dendritic tree to mimic natural odor input. (A) In each”trial” (simulation run), 25 dendritic segments (green; many are not visible in the figure because they are behind non-stimulated segments which are shown in gray) were selected randomly and identical current pulses of 5*ms* duration were injected simultaneously. A total of 200 trials were conducted. (B) Overlay of all 200 voltage traces in the proximal neurite. (C) Statistical distribution of all maximal voltage amplitudes in the proximal neurite with mean (dashed line). Error bars represent standard deviation (SD).

**Figure 3:**
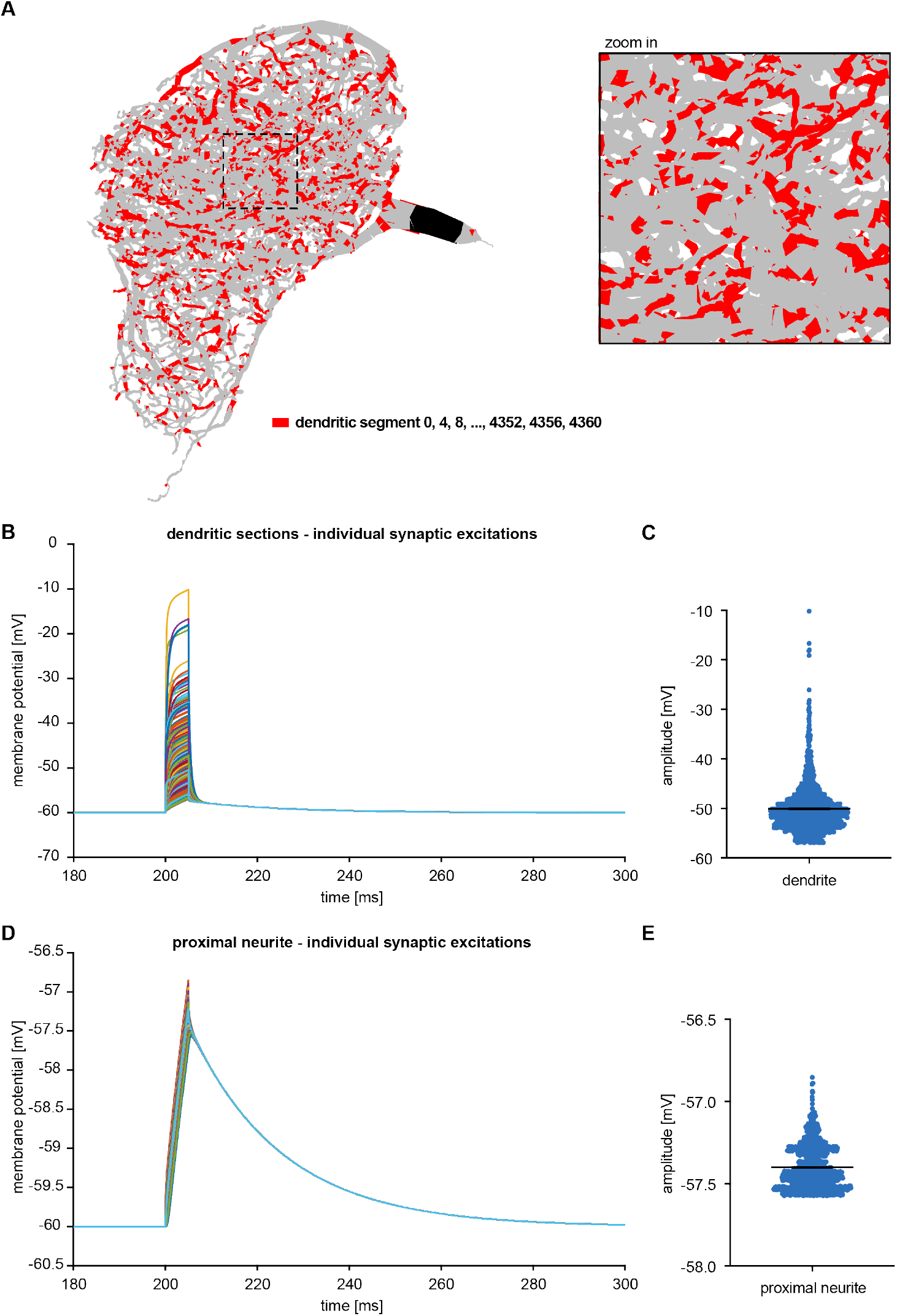
Segmentation and widespread activation of single dendritic segments to study electrotonic compactness. (A) For an even distribution every fourth segment of the dendritic tree (red) was selected for current injection. In total 1091 segments were stimulated. The voltage after current injection was measured in the stimulated segments themselves (B,C) or in the proximal neurite (black) (B,C). (B) Voltage in all activated dendritic segments. (C) Distribution of maximal voltages in all stimulated dendritic segments, with mean and SD. (D) Corresponding voltages in the proximal neurite for all activated dendritic segments. (E) Distribution of maximal voltage in the proximal neurite with mean and SD. Note the difference in vertical scales in panels C and E.

### 2.3 Distributions of voltage excursions in dendrite and proximal neurite due to individual synaptic excitations

The simulated voltage injections in Section 2.1 were applied in two randomly selected dendritic segments. Section 2.2 showed that a pattern of synaptic activation similar to what is assumed to occur in response to odor delivery to the insect reliably brings the MBON voltage above threshold. In this section we investigate the effects of current pulses injected into individual segments in the dendritic tree, and repeat this for many segments. We record the effect the injections have on the voltage of the proximal neurite as well as in the dendrite at the point of injection, and determine the voltage distributions. If the neuron is electrotonically compact, we expect the proximal neurite voltage distribution to be narrow.

Current pulses were injected systematically into segments of the dendritic tree, one at a time. We selected every fourth of the 4361 segments that NEURON divided the dendritic tree into, for a total of 1091 different segments. The segments that were stimulated are marked in red in Figure 3A. Each current pulse had a duration of 5*ms* and an amplitude of 0.02*nA*. The amplitude was selected to obtain a dendritic EPSP in approximate agreement with the observed values [8].

A plot of all evoked dendritic EPSPs is shown in Figure 3B. The statistical distribution of the maximal depolarizations (Amplitude) is plotted in Figure 3C. The average of maximal voltage excursion is —50.06*mV*, and most EPSPs fall into a range of about 10*mV*, from —45*mV* to −55*mV*. We surmise that the range of ≈ ±5*ms* for identical current injections is due to differences in local dendritic geometry.

The effect of all single synapse stimulations on the proximal neurite voltage is shown in Figure 3D and their statistical distribution in Figure 3E. The average EPSP peak at the proximal neurite was found to be −57.40*mV*. The depolarizations fall into the range [—57.57*mV*, —56.85*mV*], indicating a very compact structure. This range is narrower by an order of magnitude than the range of dendritic EPSPs (Figure 3B). It therefore appears that propagation along the dendritic structure evens out differences arising at the local dendritic site of activation, resulting in a “compactifying” effect. Equalization of synaptic input from different KCs may be of consequence for the representation of odorant information.

### 2.4 Voltage attenuation in the dendritic tree

The majority of voltages in the proximal neurite elicited by identical current injections is in a relatively small range of approximately ±5*mV* (Figure 3D). We assumed that differences in proximal neurite voltages are due at least in part to two factors.

The first factor are the differences in the distances between the stimulation sites and proximal neurite. For the selected stimulation sites (and likely for the dendritic tree as a whole), these distances range from about 2*μ*m to about 65*μ*m, see Supplementary Figure S2E,F. To test this hypothesis, we plot in Figure 4A the distance between stimulated segment and proximal neurite *vs.* the difference in maximal voltage in the proximal neurite and the dendrite. We find an overall positive correlation between these quantities, supporting our hypothesis.

**Figure 4:**
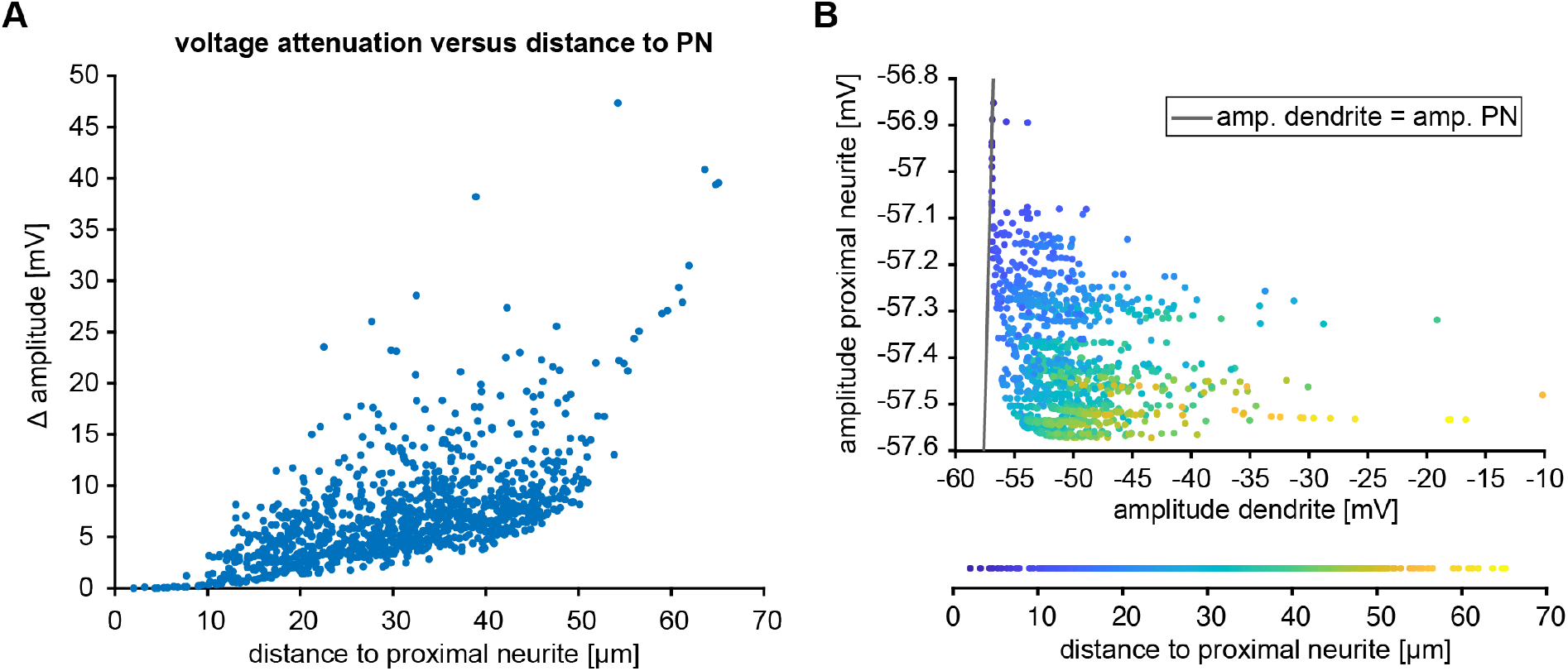
(A) Voltage attenuation between dendritic tree and proximal neurite. Attenuation is computed as the difference between the maximum voltage in each of the n=1091 stimulated dendritic segments and in the proximal neurite (Δ Amplitude). It is plotted *vs*. the distance between the dendritic segment and the proximal neurite segment. (B) Voltage excursion in the proximal neurite as a function of voltage excursion in the stimulated dendritic segment. Color code represents the distance of the stimulated dendritic section from the proximal neurite.

The second factor that we hypothesized playing a role in the voltage attenuation are the differences between dendritic voltages in response to (identical) current injection due to differences in local geometries. To understand this factor better, Figure 4B shows for each simulation the value of proximal neurite voltage as a function of the corresponding dendritic voltage. We observe a positive correlation between these two quantities, corroborating our second hypothesis. In this figure, we also use a color code to show the distance along the dendrite between proximal neurite and current injection site. This distance is generally inversely related to the proximal neurite voltage, as we already saw in Fig. 4A.

## 3 Discussion

Our numerical simulations show that the MBON-α3 neuron is electrotonically compact. As a consequence, the impact a synapse has on the output of this neuron, *i.e.* generation of action potentials, is to a large extent independent of where on its dendritic tree the synapse is located. These simulation results are in agreement with a simple analytical estimate of the space constant of this neuron. This is obtained by assuming a non-branching cylinder with the mean diameter of the dendritic segments, 0.29*μm*, as obtained from the morphological data from Takemura *et al.* [45]. We assume values for the conductance of the cytoplasm of 4.5 × 10^-5^S/cm^2^ and transmembrane conductance of 260Ω*cm*. These values were obtained by Gouwens and Wilson [18] in a combined electrophysiological and computational study of antennal lobe projection neurons, another class of *Drosophila* central neurons. The characteristic length is then computed by linear cable theory [34] as λ ≈ 350*μm*, more than three times the maximal path length between segments of the dendritic tree and the distal end of the EM reconstruction.

### 3.1 Limitations

The results obtained from these simulations are potentially limited by the available data for the specific neuron that had been modeled. While the precise morphology has recently been characterized for the MBONs in *Drosophila* [45], data describing the cell’s electrophysiological properties such as channel conductances and cytoplasmic resistivity have not been quantified. Therefore, to build the constructed model we were forced to assume that the MBON-α3 has properties similar to MBONs characterized in other insects as well as other neurons in the insect central nervous system.

Another limitation is that synaptic activation is simulated as current pulses, rather than conductance changes. This approximation is valid if interactions between synaptic currents are minimal. This may be the case here since only a very small fraction of synapses (a single one in Sections 2.1, 2.3 and 2.4, or 25 in Section 2.2, in all cases out of 12,770) is activated in the simulation. However, with the level of injected currents chosen, voltages at the proximal neurite show large deflections from rest (up to positive values) so this assumption may not be valid in all parts of the neuron. In future work we plan to go beyond this approximation, see also section 3.3.

Another potential concern is that the shape of the current injections (a square pulse) is unrealistic. It would be simple to replace it by a more realistic shape, like an α-function but it seems unlikely that this will have a substantial effect on results.

The current simulations assume identical stimulation at each synapse. As a second step it might also be interesting to use a mix of synaptic strengths from a distribution over some range. We know that KC-MBON synapses are not homogeneous entities. Cassenaer and Laurent [8] showed that the physiological impact of synapses varies over a range of at least a magnitude, and Takemura *et al.* [45] found that the physical size of synapses varies over a considerable range. The fact that we recently observed bundles of labelled CAMEL cells after paired conditioning of flies underpins this hypothesis [43]. It is possible though unproven at this time that these physiological and morphological differences in synaptic strength are caused by their plasticity during learning. The impact of such variability on the output of the MBON is of great interest. There is also variation around the mean of measured dendritic time constants, and concomitant local variations of membrane conductances. This is another target for adding a stochastic component to the simulation.

### 3.2 Deviations from physiological conditions

In our simulation experiment, we approximated the synaptic excitation pattern generated by an odor input by simultaneous current injections in 25 locations of the MBON dendrite. However, the currently published connectome gives more profound insight into definite numbers of KC-MBON connectivity. In this study, we start with a simplified excitation pattern to establish as a proof of principle. In the future, we will take into account more details in the numbers, and detailed locations, of KC-MBON synapses.

### 3.3 Model extensions

Excitability of the MBON-α3 was examined by determining whether current injections along the dendritic tree could depolarize the cell past an assumed firing threshold in the spike initiation zone (SIZ; we do not know in detail where the SIZ is located and therefore used the proximal neurite as a simple approximation for it). Future work can augment this approach in several ways. First, the input current can be modeled by replicating the dynamics of the excitatory KC–MBON synapses, beginning by incorporating into the model representations of spiking KCs releasing excitatory neurotransmitter onto receptors within postsynaptic densities located at identified KC-MBON synapses on the MBON dendrites. Additionally, active channels can be inserted into the membrane of the putative SIZ to reproduce spiking behavior of the MBON-α3. Active channels may also be potentially distributed along the neuron’s dendrites as a potential means of aiding the propagation of current from the synapse locations to the SIZ. These channels would only strengthen the depolarization at the SIZ, so the determination that stimulation of KCs can sufficiently depolarize the SIZ past a threshold is unlikely to be altered. Incorporation of these active channels would require that their conductances and dynamics be characterized in Drosophila in order for the model to be effectively parameterized. Future work can further explore associative olfactory learning in insects by augmenting the model constructed in this study to incorporate additional elements of the mushroom body such as the effect of dopaminergic feedback on the KC-MBON synapses.

## 4 Materials and methods

### 4.1 Computational Model

Morphology data for the Drosophila MBON-α3 was originally characterized using scanning electron microscopy [45] and for the present study obtained from the database NeuroMorpho.org [1]. These data describe the structure of the its dendritic arborization throughout the mushroom body output neuron α3 compartment (Figure 1B, S1A). The reconstruction is limited to the portion of the neuron in the mushroom body, not including its soma and axon.

Electrophysiological properties of the neuron were determined through a literature search, as noted below. In the absence of data specific to the *Drosophila* MBON-*α*3 neuron, electrophysiological parameters were chosen based on the properties of MBON neurons in other organisms (*e.g.*, locusts) considered alongside values that would be typical throughout the Drosophila central nervous system (Table 2). For the question of electrotonic compactness, the resting (passive) transmembrane conductance is obviously crucial. We were not able to find data that reported its value explicitly for MBON neurons. Therefore we estimated its value from two sources. One is the measured dendritic time constant of *β*-Lobe Neurons (*β*LNs) recorded by Cassenaer and Laurent [8]; their Figure 1b. Note however that (a) this is a different cell class (not *α*-MBONs) (b) in a different insect (locust not Drosophila), that (c) the set of recorded neurons is small (*N* = 9) and that (d) the value is an average but the variation around this average is substantial. Despite all these limitations, the values chosen are in a physiologically plausible range and we feel that they are the best estimate we can use given the extant data. The other data point is the specific membrane capacitance which is assumed to take the value *C_m_* ≈ 1*μF/cm*^2^ based on the biophysics of cell membranes [33], in excellent agreement with measured values [17].

**Table 2:**
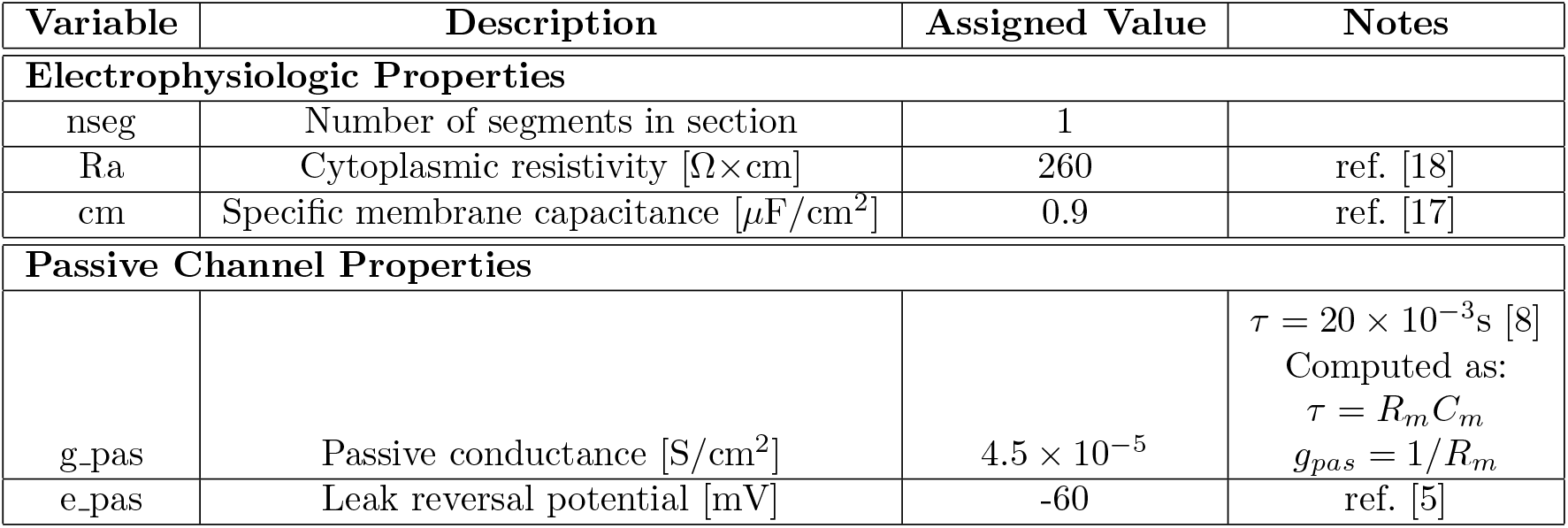
Electrophysiological properties of the Drosophila MBON-*α*3 used in the NEURON environment. Parameter values are obtained either directly through literature search or, where necessary, calculated based on available data, as noted. The passive conductance *g_pas_* was computed from the cell time constant [8] and the specific membrane capacitance *C_m_*, see Section 4.1 for details.

All simulations were performed using the NEURON 7.6.2 simulation environment[20] running through a Python 2.7.15 interface in Ubuntu 18.04.1. The morphology data divided the MBON-α3 dendritic tree into 4361 small sections averaging 1.03*μm* in length and 0.29*μ*m in diameter. Each section was automatically assigned an index number by NEURON. Each of these sections was modeled as a piece of cylindrical cable, with connectivity specified by the neuron’s morphology. Passive leak channels were inserted along the membrane of the dendrite sections (Table 2). To determine electrotonic compactness, current was injected into selected dendritic compartments and their effect at the proximal neurite studied. Resting potential is always assumed as −60mV. Membrane potential data was analysed and plotted with Matlab (MathWorks) and Prism (GraphPad).

### 4.2 Immunohistochemistry

Flies were reared on standard fly food at 25°C and 65% humidity. Male flies expressing MB082C-Gal4 [2] and 10xUAS-mCD8GFP (BDSC 32186 [38]) were fixed for 3.5h at 4°C in 4% PFA containing PBST (0.2% Triton-X100). Flies were then washed for at least 3 x 30 min at RT in PBST before dissection. Dissected brains were then labelled with primary antibodies (rabbit anti-GFP (A6455, Life technologies, 1:2000) and mouse anti-Brp nc82, Developmental Studies Hybridoma Bank, Iowa, 1:200) for two nights at 4°C. After incubation, brains were washed at least 6 x 30 min in PBST. Secondary antibodies (Alexa488 (goatαrabbit) and Alexa568 (goatαmouse) coupled antibodies, Life technologies, 1:1000) were applied for two nights at 4°C. After a repeated washing period, brains were submerged in Vectashield (Vector Laboratories), mounted onto microscope slides and stored at 4°C.

Images were acquired using a confocal scanning microscope (Zeiss LSM 710) and a 25x (Zeiss Plan-NEOFLUAR, NA 0.8 Korr DIC) oil objective. Raw images were processed with Fiji [41] and Adobe Photoshop. Uniform adjustments of brightness and contrast were performed in fiji and Adobe Photoshop.

## Acknowledgments

Work supported by NSF 1835202, NIH R01DA040990 and NIH R01EY027544.

## Supplementary Material

**Figure S1:**
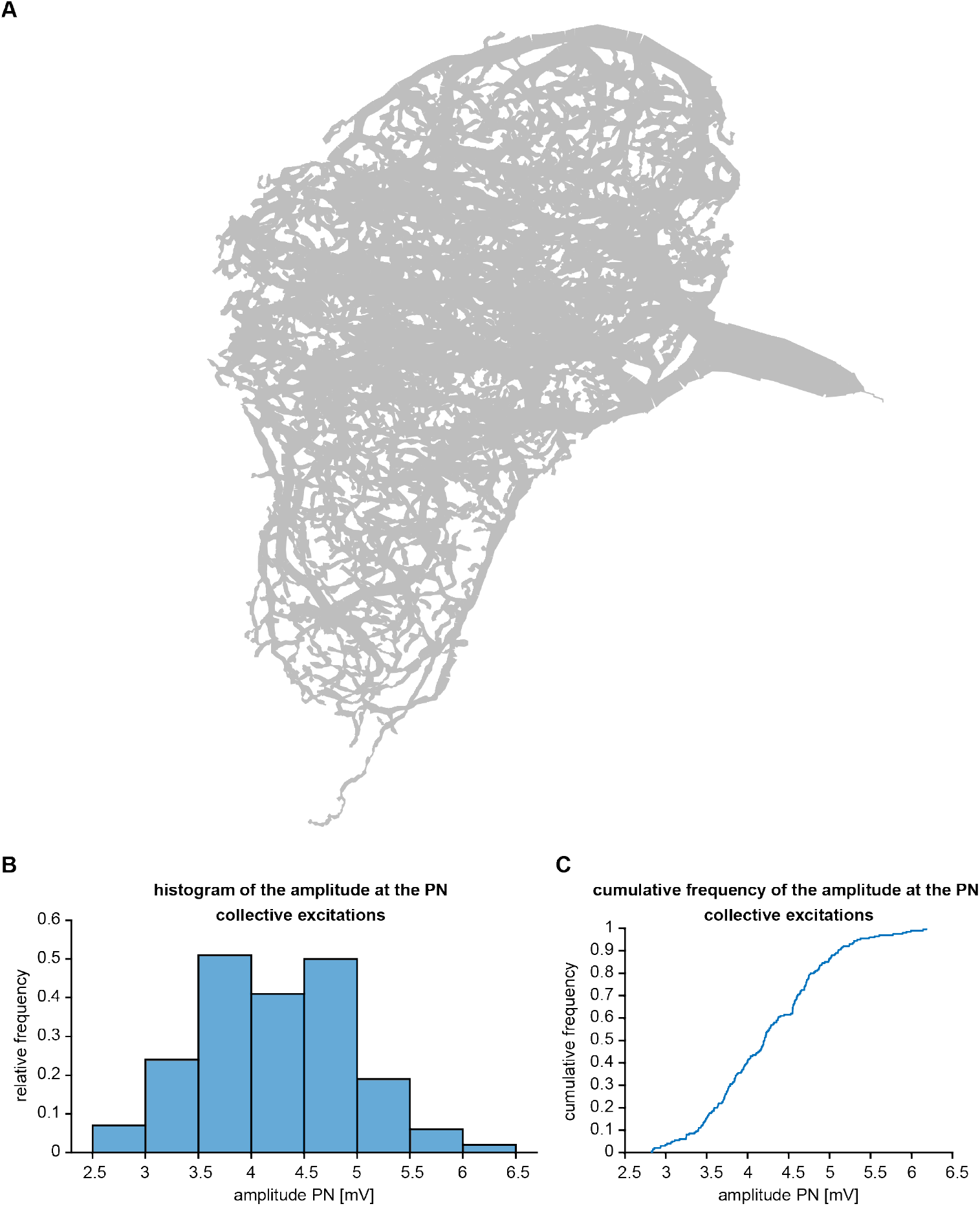
Projection of dendritic morphology and statistics of the amplitudes from collective excitations. Amplitudes were measured through 200 simulation runs of collective excitation patterns, each consisting of current injection into 25 randomly selected dendritic segments. (A) Projection of the complete dendritic reconstruction morphology. (B) Relative frequency distribution of the amplitude from dendritic stimulation for all 200 excitation patterns. Bin width 0.5*mV*. (C) Same, but shown is the cumulative frequency distribution.

**Figure S2:**
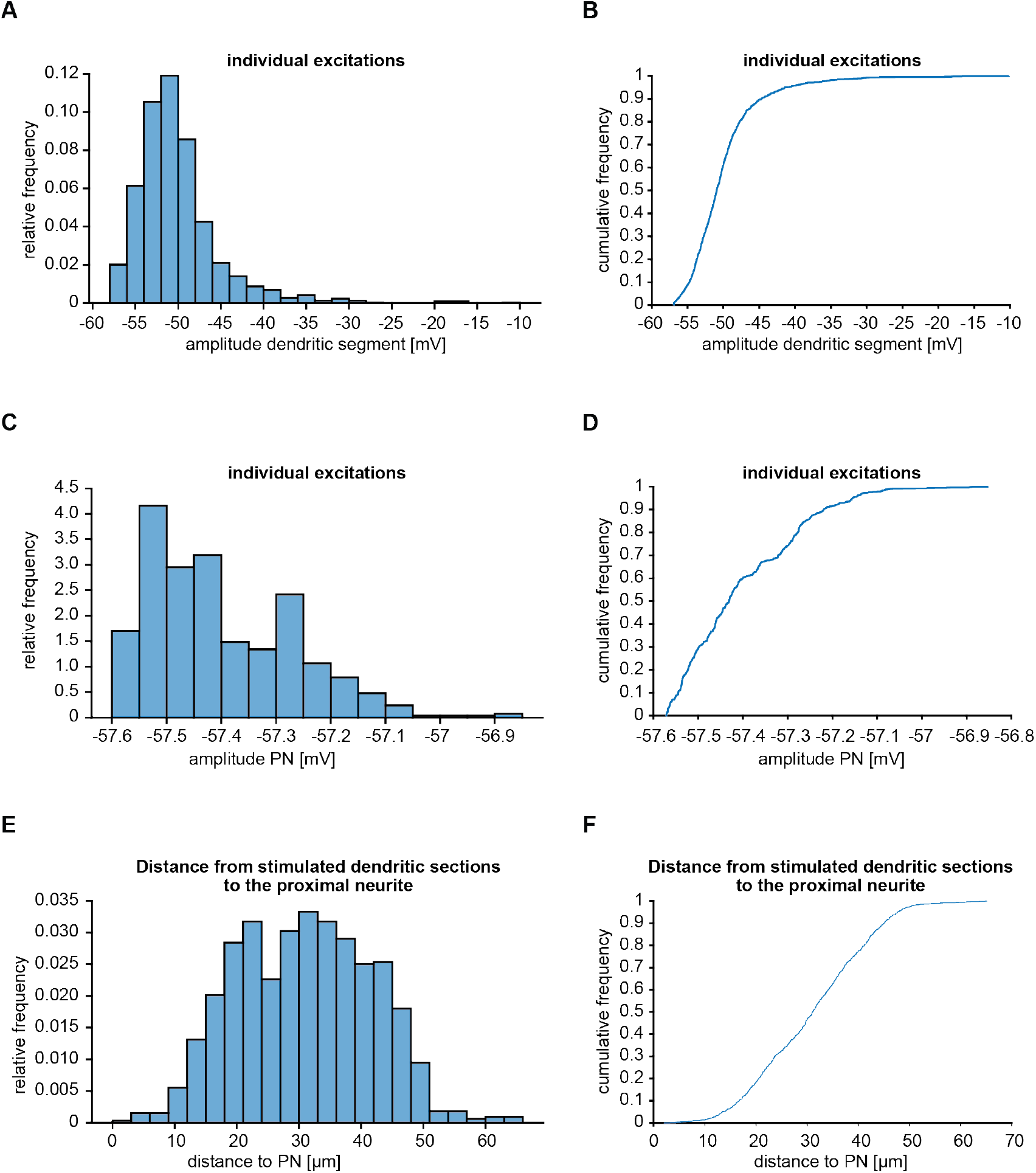
Distribution of maximal voltages in dendritic tree and proximal neurite, demonstrating the electrotonic compactness of the dendritic tree. Voltage excursions are responses to the injection of identical currents in 1091 randomly selected segments in the dendritic tree (A) Relative frequency distribution of maximal voltage amplitude in stimulated dendritic segments. Bin width *2mV*. (B) Same, but shown as cumulative frequency distribution. (C) Same as (A) but measured in the proximal neurite. Bin width 0.05*mV*. (D) Same as (B), but measured in the proximal neurite. (E) Relative frequency of the distance along the dendrite of a stimulated dendritic segment from the proximal neurite. Bin width 3*μm* (F) Same, but shown is the cumulative frequency distribution.

## Footnotes

1 We chose to make the current injection very long (100ms) to demonstrate the nearly perfect overlap of the proximal neurite voltages for different stimulated segments (Figure 1H) even for this long duration. As mentioned, the current amplitude was adjusted to result in a realistic dendritic EPSP size.

